# Precise Regulation and Site-specifically Covalent Labeling of NSUN2 Enable by Genetic Encoding Expansion

**DOI:** 10.1101/2020.12.22.424046

**Authors:** Jizhong Zhao, Hongmei Hu, Sheng Wang, Li Wang, Rui Wang

## Abstract

RNA plays a critical role in gene expression regulation, cell migration, differentiation, cell death in living organism. 5-Methylcytosine is a post transcriptional RNA modification identified across wide ranges of RNA species including message RNAs. It is reported the addition of m^5^C to RNA cytosines is enabled by use of NSUN family enzyme, NSUN2 is identified as a critical RNA methyltransferase for adding m^5^C to mRNA. We demonstrated here that natural lysines modified with special groups were synthesized via chemical synthesis. Through two rounds of positive screening and one round of negative screening, MbPylRS-tRNAcua unnatural lysine substitution system which can specifically recognize lysine with defined group was evaluated and identified. Non-natural lysine substitution at C271 of NSUN2 active site and subsequently fluorescent labeling was realized via so-called click reaction. The function of NSUN2 mutant and its upregulated CDK1 gene and its effect on cell proliferation were also evaluated.

---

Ribonucleic acids play critical roles in gene expression regulation, cell migration and differentiation, cell death in living organisms. ^1–2^ In addition to the four canonical nucleotides, there are over 150 chemical modifications in endogenous nucleic acids enable their diversified structures and function. ^3–5^ To achieve the regulation purpose, RNA modifying enzymes served as essential roles, that are responsible for adding wide range chemical modifications into target RNAs. Methylation has been found to be heavily attached into intrinsic RNAs across wide scopes of species. ^6–12^ Besides the most heavily modification termed m^6^A in mammalian animal, the m^5^C modification (abbreviated as m^5^C, **Figure 1A**) enable by varied enzymes, i.e., NOL1/ NOP2/ SUN domain (NSUN) family, has attracted broad attentions in recent years. ^13–15^ It is known that 5-methylcytosine is a post transcriptional RNA modification identified across wide ranges of RNA species including message RNAs. It is reported that the addition of m^5^C to RNA cytosines is enabled by use of NSUN family enzymes as well as the DNA methyltransferase DNMT2 in mammalian cells. NSUN2 is identified as a critical RNA methyltransferase for adding m^5^C to mRNA. Most recently, Yang and coworkers have revealed that m^5^C modification was enriched in CG-rich motif. ^14–15^ These regulated regions immediately downstream of translation initiation sites and has conserved features across mammalian transcriptomes. Moreover, it was disclosed that m^5^C is recognized by the mRNA export adaptor ALYREF as shown specifically by *in vitro* and *in vivo* studies, which NSUN2 modulates ALYREF’s nuclear-cytoplasmic shuttling, RNAbinding affinity and associated mRNA export. ^13^ Regarding these crucial roles of NSUN family enzyme in m^5^C-involving RNA biological activities in living organisms, namely cellular proliferation/senescence/migration/differentiation, mRNA nuclear export, enhanced mRNA translation, tRNA stabilization and cleavage, ^16^ to achieve the site-specific modulation of the NSUN enzyme is therefore of high importance. Regrettably, precise regulation of the nuclear-cyttoplasmic shuttling of endogenous RNAs via manipulation of the activity of NSUN family enzyme (i.e., NSUN2), the current approach remains elusive.

**Figure 1.**
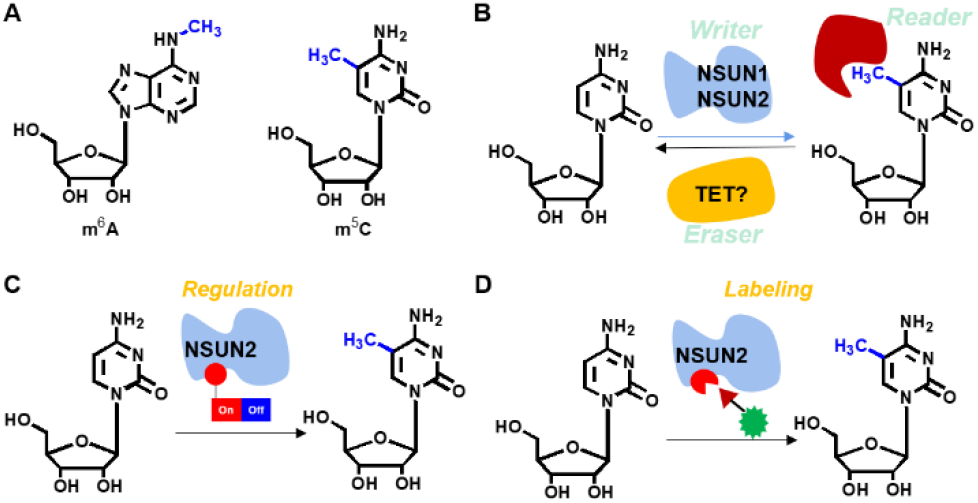
RNA m^5^C modification and its invoked enzymes and regulation and labeling strategy. (A) Representative modifications in RNA such as m6A, and m^5^C etc. (B) m^5^C involved enzyme, including the “writer” enzyme NSUN2. (C) Our strategy of a switch precisely setting on the NSUN2 that regulate the RNA methyl modification on cytosine 5 position by a gene encoding expansion technique. (D) Site-specific labeling could be achieved by further bioorthogonal reaction.

In another hand, the genetic encoding expansion technique has been pioneered by Schultz and coworkers and demonstrated successfully in the past decades, serving as a powerful tool in molecular biology such as identification of PPIs, regulation of proteins/ RNAs, drug discoveries *etc*.^17–22^ Specifically, tremendous achievements have been made in protein regulation in recent years, for instance., Chen and workers have utilized genetic encoding expansion technique to perturb and manipulate the functions of various enzymes such as FTO, luciferase and KRAS.^22^ Based on one of our previous studies, ^23^ It was demonstrated that we could install the non-canonical PBBK into the critical site of Cas9 endonuclease through genetic encoding expansion, thus the precise regulation of CRISPR-Cas9 gene editing have been achieved. We therefore speculated that the NSUN2 is also able to be regulated via the genetic encoding expansion technique. Moreover, if the defined functionality could be installed at the same time, the regulation of m^5^C level on specific RNAs in spatiotemporal manner and further site-specific labeling of NUSN2 could also be achieved. As a consequence, the intrinsic nature of m^5^C modifying enzyme NSUN2 could be further evaluated and testified, and the upregulated genes could further be clarified.

To illustrate the feasibility of our method, we synthesized the four lysines substrates in advance. As depicted in **Scheme 1**, the four lysine substrates were constructed, and all data were identified by corresponding NMR and mass spectrum (**Scheme 1A**). Typically, **S3** were obtained in linear two steps from commercially available Boc-protected *L*lysine in over 95% yield (**Scheme 1B**). Notably, all of the four lysine analogues were able to be transformed into their native forms by the established methods. ^24–26, 23^ For instance, substrate **S1** has been demonstrated to be restored into lysine by UV (365 nm)-light in 5 minutes. ^24^ Substrate **S2** could be transformed to its untreated form by basic hydrolysis. ^25^ Substrate **S4** could also be rescued into lysine by signaling molecule hydroperoxide as reported. ^23^ Notably, we demonstrated herein that substrates **S3** was able to be restored to its native form by addition of TCEP (tricarboxyethyl phosphine) in a relatively rapid manner, as reported before. ^26^ Kinetics investigation of the reaction of **S3** with TCEP displayed a pseudo first-order with half time to be 24.8 minutes (**Scheme 1C, 1D**).

## Screening of site-specifical unnatural amino acid substitution system

We next turned to illustrate whether our substrates (**S1-S4**) could be site-specifically incorporated into enzyme NSUN2. Regarding the structure feature of m^5^C modification enzyme NSUN6,^27^ it was reported that the cytosine on 271 position was essential, playing crucial role in the process of RNA m^5^C modifications. Unfortunately, the crystal structure of NSUN2 is unknown. By screening NSUN2 mutant incorporated with our lysine analogues (i.e., our substrates **S1-S4**) randomly, the regulations of NSUN2 enzyme activity could be achieved. Regarding for the specific substrate **S3**, the terminal 6-amino group has been transformed to azido group comparing with lysine. As a consequence, the NSUN2 activity could be restored if the Staudinger reaction was applied to reduce azido group to amine functionality. ^27^ Moreover, the bioorthogonal reaction of azido group with various alkynes (i.e., attached with fluorescence or biotinated functionality) could also be utilized, therefore the fluorescent labeling or chemical pulldown could be realized.

Next, we focused on screening of tRNA transferase (MbPylRS) that specifically recognizes lysine derivatives with defined modified groups.

## Screening of specific MbPylRS

Firstly, the mutation library was obtained via random mutation of MbPylRS active site. In the context, random mutations were carried out in six sites of MbPylRS, namely L266, L270, Y271, L274, C313 and W383, and 10^8^ mutants were theoretically produced. MbPylRS mutants which can specifically recognize lysine derivatives (**S1-S4**) were screened from the mutant library. To facilitate screening process, we constructed two screening systems.^18–20^ The first screening system was positive screening system, and the second screening system was negative screening system. Based on our two screening systems, MbPylRS-tRNAcua system with specific recognition and coding ability for target lysine derivatives (**S1-S4**) has been identified through two rounds of positive screening and one round of invisible screening. Relying on this system, the efficient replacements of unnatural lysines (**S1-S4**) at NSUN2 essential sites have achieved, as shown in Supplementary **Figure S21**.

## Construction of dual fluorescence report system for detection of unnatural lysine substitution system

To evaluate whether the screening of unnatural lysine substitution system is feasible, a dual fluorescence reporter system was constructed in advance. HeLa cells expressing MbPylRStRNAcua and dual fluorescent reporter gene were cultured in DMEM medium containing specific lysine derivatives (**S1-S4**). The expression of fluorescent protein was detected after 24 hours. The results showed that MbPylRS-tRNAcua, which could specifically recognize the four lysine derivatives (**S1-S4**), has successfully been selected after three rounds of screening, and the unnatural lysine substitution of NSUN2 in eukaryotic cells was successfully realized. As shown in **Scheme 2B and 2C**, it revealed that the expression of red fluorescent protein cannot be detected in the absence of **S1-S4**, while the expression of two fluorescent proteins can be detected in the presence of **S1-S4**, indicating that the MbPylRS-tRNAcua identified above was able to realize the recognition of specific non-natural lysine and code the amber codon, and efficient replacement of lysine with defined **S1-S4** on NSUN2 was achieved.

## Substitution with unnatural amino acids at the essential site of NSUN2

Next, we turned to explore the feasibility of substitution with the unnatural lysine on essential site of NSUN2. It has been demonstrated that C271 was a critical site of NSUN4 (Supplementary **Fig. S17**), which was responsible for the separation of NSUN from substrate after catalytic activity completion. ^13–15^ We thus focus on construction of C271 NSUN2 mutant in eukaryotic cells. The eukaryotic expression system of NSUN2 was firstly constructed, regarding that the C271 mutation of NSUN2 gene sequence is TAG amber codon, and the C-terminal of NSUN2 labeled with EGFP fluorescent protein. In MbPylRS-tRNAcua expression cells, NSUN2-EGFP eukaryotic expression system was transiently stained. The cells were cultured in DMEM medium containing varied lysine derivatives (**S1-S4**). After 24 hours, the cells were lysed and the total protein was extracted, the protein expression of NSUN2 was further detected by western blot assays. The results demonstrated that the expression of NSUN2 C271 mutant protein could be detected in the presence of **S1-S4** (**Figure 2B & 2C**). The expression level of NSUN2 protein in the presence of **S1** and **S3** was relatively high (**Figure 2B)**, while the NSUN2 protein in the presence of **S2** and **S4** were also detectable, but much lower that **S1** or **S3** substrates. Interestingly, in addition to the target expression, there was a 2.0 KD band have also been expressed and observed, which may due to the cotransfection process.

## Detection of NSUN2 using S3 (lys-N3) by dual fluorescent reporter system in eukaryotic cells

To investigate whether the NSUN2 could be subcellularly located and detected, we turned to the click reaction considering the azido functionality on NSUN2. It was demonstrated that in the presence of **S3**, the unnatural lysine substitution of NSUN2 mutation C271 could be realized efficiently. Next, NSUN2 with **S3** incorporated on C271 site was labeled with DBCO-Cy5 fluorescent dye through click reaction. After labeling, we clearly observed the subcellular location of the NSUN2 mutant (**Scheme 2D**). To further evaluate the accuracy of this protein labeling method, the EGFP fusion protein of NSUN2 was constructed (**Scheme 2D**). It was demonstrated that the subcellular localization of NSUN2 was detected through monitoring of the co-localization of DBCO-Cy5 (red fluorescence) and EGFP (green fluorescence) (**Scheme 2D**). In addition, in order to improve the expression efficiency of NSUN2, the concentration of **S3** was further optimized. The cells were cultured in DMEM medium containing 1.0 mM, 2.0 mM, and 3.0 mM of **S3** respectively, and the protein localization was detected by fluorescence microscope after staining with DBCO-Cy5 (Supplementary **Scheme S5 and S6**). The results revealed that with the elevated concentration of **S3**(lys-N3), the substitution efficiency of unnatural lysine in the process of NSUN2 protein expression enhanced. Through the detection of localization of green fluorescent protein and red fluorescence, it was obviously observed that the dual fluorescence was co-located, which indicated that **S3** has been efficiently incorporated at NSUN2 C271 site, displaying potential to enable real-time tracking of NSUN2. Our result demonstrated that the biological orthogonal reaction of unnatural lysine with azido group together with the genetic encoding expansion enabled the specific NSUN2 protein labeling and tracking **(Scheme 2C and 2D)**.

## Effect of NSUN2 active site mutation on function

To further evaluate the effect of NSUN2 C271 mutation on the activity of NSUN2, we tried to explore mechanism of its upregulated gene. Firstly, the effect of NSUN2 C271 mutation to alanine was evaluated. Serving as an RNA methyltransferase, the mutant on NSUN2 perturb the RNA methylation would further affect the function and stability of target RNA. Among them, it was found CDK1 has been identified to be the upregulated gene of NSUN2. ^28–30^ It was reported that NSUN2 was highly expressed in a variety of tumor cells, and the expression of NSUN2 promoted the translation process of CDK1, thus promoting the process of cell cycle and cell proliferation. ^30–32^ In this context, we therefore knocked out the expression of NSUN2 in HeLa cells, and then replenish the wild-type or C271 mutant of NSUN2. Subsequently, the proliferation ability of HeLa cells was detected by CCK8 kit, and the effect of NSUN2 on CDK1 transcription level was detected by RT-PCR. The results showed that NSUN2 mutant had no significant effect on the transcription level of CDK1, but the mRNA level of CDK1 resulted in a 50% decrease after the NSUN2 mutation, as shown in **Scheme 3A**. However, when NSUN2 was knocked out, the cell proliferation level decreased significantly, and when the cell was supplemented with wild-type NSUN2, the cell proliferation level returned to HeLa wild-type level. ^32^ Notably, when NSUN2 C271 was mutated to alanine, the cell proliferation was significantly reduced. Thus, it was demonstrated that the C271 mutation significantly inhibited the activity of NSUN2 (**Scheme 3B**).

## Effect of unnatural lysine substitution on the function of NSUN2

Next, the regulation of NSUN2 when C271 essential site replaced by unnatural lysine (**S1-S4**) were further investigated. In HeLa cells, MbPylRS-tRNA_cua_ unnatural lysine substitution system and NSUN2-C271TAG mutant were co expressed in HeLa cells. The cells were cultured in DMEM medium containing **S1-S4**, and the proliferation ability of tumor cells was detected. The results showed that the cell proliferation level in **S1** and **S3** medium was significantly lower than that in wild type, and was significantly higher than that in **S2** and **S4**. It thus hinted that **S1** and **S3** displayed higher replacement efficient than **S2** and **S4** at NSUN2 C271 sites (**Scheme 3C**).

As an m^5^C methyltransferase, the studies of function of NSUN2 is still in its infancy stage, especially the regulation on its RNA substrate.^33^ The regulation of essential sites of NSUN2 is an efficient approach to manipulate the function of this enzymes. It was reported that there are mainly two active sites of NSUN2, namely C321 and C271, in which C321 is responsible for the methyl transfer and C271 is responsible for the completion of cytosine methylation, and NSUN2 is separated from the modified site of cytosine. Through genetic encoding expansion approach, to prevent the perturbation of the catalytic activity of NSUN2, we focus on modification of the C271 site of NSUN2 with varied lysines, and subsequently evaluated the effect of NSUN2 C271 mutation on its activity and upregulated gene. We demonstrated that lysines with four different modifications have been successfully added at C271 site of NSUN2 by genetic code expansion technique. In addition, RT-PCR and cell proliferation assay showed that C271 mutation had minimal effect on the mRNA level of upregulated factor CDK1, but significantly inhibited the function of CDK1. C271 mutation with **S1-S4** could inhibit the proliferation of HeLa cells. This may be due to the C271 site mutation, which makes NSUN2 unable to separate from the target RNA sequence after completing the catalysis of m^5^C, thus affecting the function of RNA, such as the translation process of target protein CDK1, ^28^ thus inhibiting the proliferation of cells.

In summary, we demonstrated here that natural lysines modified with special groups were synthesized by organic synthesis. Through two rounds of positive screening and one round of negative screening, MbPylRS-tRNAcua unnatural lysine substitution system which can specifically recognize lysine with specific group was evaluated and identified. Non-natural lysine substitution at C271 of NSUN2 active site and subsequently fluorescent labeling was realized via the so-called click reaction. The function of NSUN2 and its upregulated CDK1 gene and its effect on cell proliferation were also evaluated. Further clarifying of regulation NSUN2 is under way on my lab and will be reported in due course.

In a word, by employing genetic encoding expansion, we successfully constructed the NSUN2 model with essential site mutated with unnatural lysine bearing N3, PBin, NO2, CF3 etc. Relying on the click reaction, we fulfilled the efficient labeling and regulation of NSUN2, which lays a foundation for further research on the function and regulation mechanism of upregulated gene.

## ASSOCIATED CONTENT

The Supporting Information is available free of charge on the ACS Publications website. brief description (file type, i.e., PDF).

## AUTHOR INFORMATION

### Author Contributions

### Notes

The authors declare no competing financial interests.

## ACKNOWLEDGMENT

R. Wang thanks the starting fund from Huazhong University of Science and Technology for financial support. One-Hundred-Talents Youth Program of Hubei Province is appreciated. Jason Chin is high appreciated for generous gift of the plasmid of *Pbk-pYlS, Prep-PylT, pBar-PylT*. We thank Prof. Chaomei Xiong for NMR experiment and Prof. Lingkui Meng for mass spectrum.

## ABBREVIATIONS

CDK: cyclin-dependent kinases
NSUN2: NOP2/Sun RNA methyltransferase 2
DBCO-Cy5: DBCO-Sulfo-Cy5
FTO: fat mass and obesity
KRAS: Kirsten rat sarcoma viral oncogene homolog

---

**Scheme 1.**
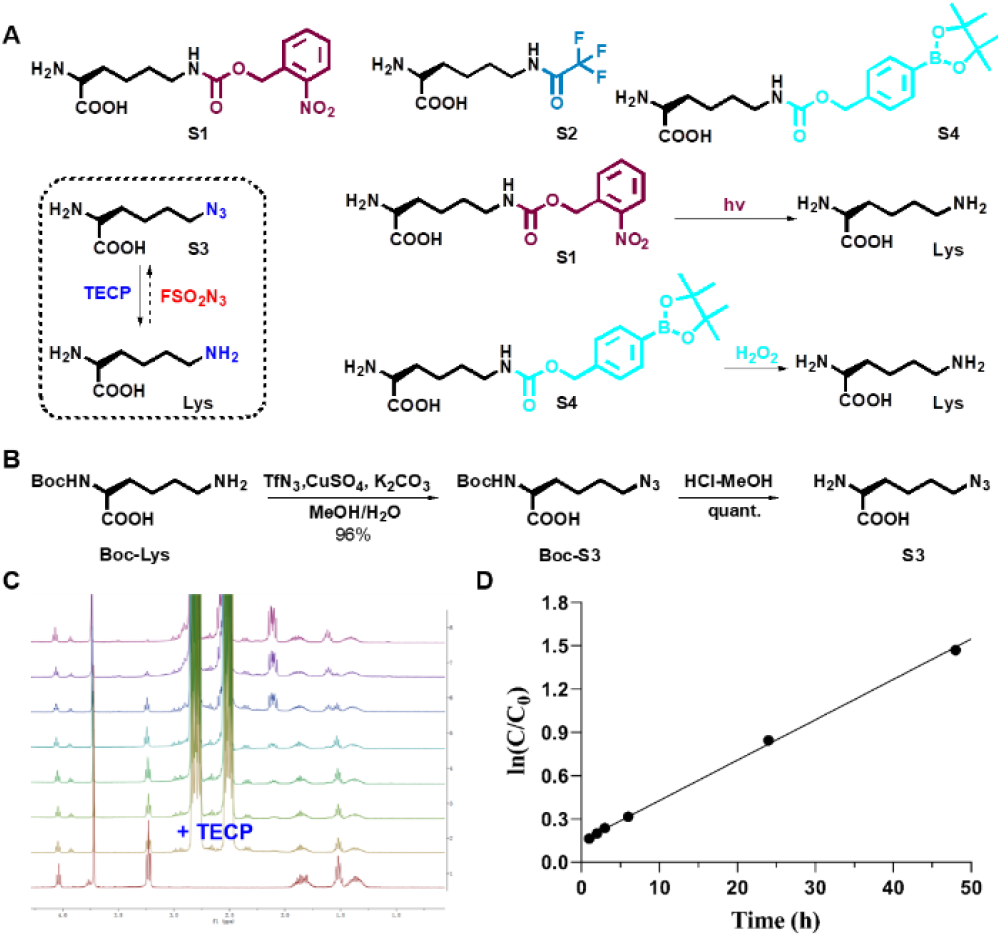
Investigations of the lysine analogues in our studies. (A) Substrates (**S1-S4**) for gene encoding expansion and their reversible methods triggered by small molecules or UV-light. (B) Synthetic approach for 6-azido lysine (**S3**). (C) Dynamic investigation of Staudinger ligation of **S3** with TCEP (10 equivalents, in D2O) by use of in-suite proton NMR experiment. (D) Reaction order plot. It was anticipated the pseudo-first order reaction of **S3**(20 mM) with TCEP (200 mM) in D2O were observed, with slope 0.02798, half time t1/2> 24.8 min. TCEP > Tris (2-carboxyethyl)-phosphine.

**Scheme 2.**
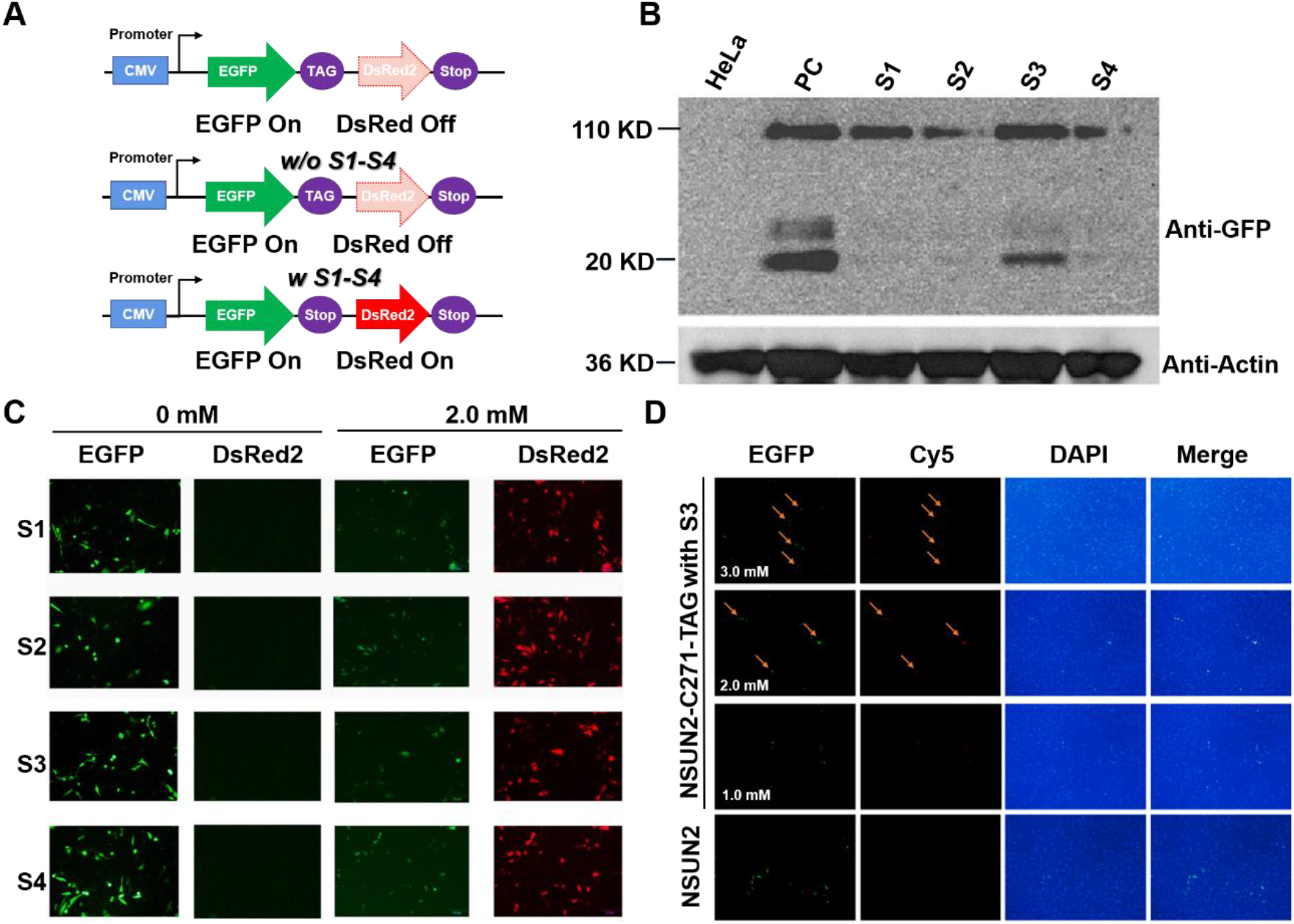
Constructions of dual reporter system for NSUN2. (A) Construction of dual fluorescent reporter system. (B) Western blot was used to detect the effect of unnatural amino acid substitution on the expression of NSUN2 protein. HeLa cells expressing MbPylRS-tRNAcua system were cultured in DMEM medium containing **S1-S4**(2.0 mM) synthetic lysine. After 24 hours, the cells were lysed and the total protein was extracted. The protein expression was detected by Western blot. HeLa cells transfected with NSUN2 overexpression plasmid were used as positive control group. (C) Detection of specific sites of unnatural and unnatural amino acid insertion proteins by dual fluorescence reporting system. (D) HeLa cells expressing MbPylRS-tRNAcua amino acid substitution system and GFP-NSUN2 C271TAG mutation system were cultured in DMEM medium containing different concentrations of **S3**(1.0, 2.0, and 3.0 mM) for 24 hours, and then DBCO-Cy5 (50 μM) was used to react with lys-N3 to label NSUN2. NSUN2 was labeled with GFP, and the subcellular localization of NSUN2 can be detected by fluorescence microscopy.

**Scheme 3.**
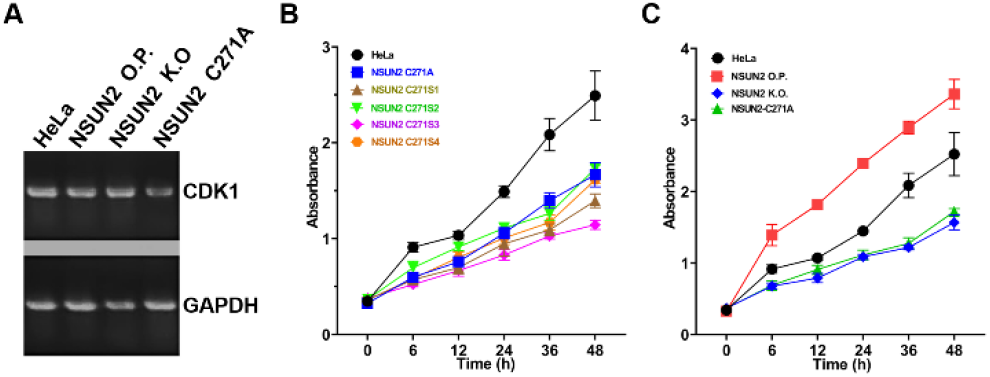
Investigations of NSUN2 mutant towards cell activity. (A) Fig. 3A effect of NSUN2 active site mutation on downstream gene CDK1 RNA transcription and cell proliferation. The A map was used to detect the effects of NSUN2 overexpression, knockout, and C271A on the transcriptional level of the downstream gene CDK1 in HeLa cells, and the mRNA level of CDK1 was detected by RT-PCR agarose gel electrophoresis. B diagram shows the effect of NSUN2 overexpression, knockout and C271A mutation on cell proliferation. (B) It effects of unnatural amino acid substitution at NSUN2 active site on cell proliferation after NSUN2 C271 was replaced by **S1-S4** in HeLa cells, CCK8 was used to detect cell proliferation.

